# Characterization of a novel R98Q mutation that confers resistance to sulfentrazone in common ragweed (*Ambrosia artemisiifolia*) populations from Michigan

**DOI:** 10.64898/2026.07.31.742095

**Authors:** Sara Álvarez-Rodríguez, Michael Ozolins, Aimone Porri, Jens Lerchl, Eric Patterson

## Abstract

**BACKGROUND:** In 2023, soybean growers in Eaton County, Michigan, reported repeated failures to control common ragweed (*Ambrosia artemisiifolia* L.) with (PPO)-inhibiting herbicides, sulfentrazone and fomesafen, in non-genetically modified soybeans. This study aimed to investigate resistance levels and mechanisms of resistance in two suspected resistant populations (R1 and R2).

**RESULTS:** Dose–response assays revealed resistance to sulfentrazone in both populations, with LD_50_ values 24 and 36-fold higher than the susceptible population and reduced sensitivity to fomesafen. Nanopore sequencing identified two PPO2 target-site substitutions at codon 98: Arg- 98-Leu (R98L) and a novel Arg-98-Gln (R98Q) mutation. Both substitutions were associated with reduced herbicide binding affinity in computational modelling simulations. R98Q substitution conferred strong and selective resistance in the PPO2 enzyme inhibition assays, supporting its role as a key resistance mechanism.

**CONCLUSION:** This study reports the first field occurrence of the R98Q substitution in PPO2 of *A. artemisiifolia* populations and demonstrates its association with to PPO inhibitor resistance. These findings highlight the rapid evolution of target-site resistance and the need for continued monitoring and rapid diagnostic tools to detect emerging mutations. Further research is needed to clarify the contribution of additional resistance mechanisms.

## 1. INTRODUCTION

*Ambrosia artemisiifolia* L., known as common ragweed, is one of the most problematic weed species within the Asteraceae family. While native to North America; its strong adaptive capacity to diverse environments, high reproductive potential, and prolific pollen production have facilitated its global spread over the last century. As a result, *A. artemisiifolia* has become an invasive species worldwide, with established populations across Europe and Asia^1,2^. This species poses a serious threat to crop yield, having evolved resistance to herbicides representing at least five distinct sites of action (SOA), including chemistries from HRAC Groups 2, 4, 5, 9, and 14^3^. In cropping systems, even low-density infestations can cause substantial yield losses. For example, Hall et al. (2021) reported that a single *A. artemisiifolia* plant per square meter reduced soybean [*Glycine max* (L.) Merr.] nodulation by 56%, resulting in an 18% yield reduction. Beyond its agronomic impact, *A. artemisiifolia* is also a major public health concern due to its highly allergenic pollen, which is a leading cause of hay fever, allergic rhinitis, conjunctivitis, and asthma in sensitized individuals^5^. Because of its invasive spread, its impact on crop yields, and its growing herbicide resistance, *A. artemisiifolia* is difficult to manage and requires effective control strategies in cropping systems.

Group 14 herbicides target the protoporphyrinogen IX oxidase (PPO) enzyme, which catalyzes the oxidation of protoporphyrinogen IX to protoporphyrin IX, a key precursor in chlorophyll and heme biosynthesis^6^. Resistance to PPO-inhibiting herbicides has been increasingly reported, with at least 18 weed species worldwide evolving resistance to this mode of action^3^. Target-site resistance to PPO-inhibiting herbicides is primarily associated with mutations in the *PPO2* gene. Among these, substitutions at residue Arg128 (R128) have been widely reported in amaranth species, particularly in Palmer amaranth (*Amaranthus palmeri* S. Watson) and waterhemp [*Amaranthus tuberculatus* (Moq.) J.D. Sauer], including R128G, R128M, and R128L ^7,8^. Similar substitutions at this position have also been reported in other species, such as wild poinsettia (*Euphorbia heterophylla* L.), where the R128L mutation conferred cross-resistance to multiple PPO-inhibiting herbicides^9^. This residue corresponds to Arg98 in *A. artemisiifolia* and represents a key position involved in PPO-inhibitor resistance across species^10^. In addition, a codon deletion at position 210 (ΔG210) has been widely documented in amaranth species^11^ while a substitution at Gly399 (G399) has also been associated with resistance to PPO inhibitors in *A. palmeri*^12^.

Together, these mutations highlight the importance of target-site alterations in the evolution of resistance to PPO-inhibiting herbicides.

PPO-inhibiting herbicides are a critical control tool for weeds in non-GM soybean and dry bean production, as they do not have access to broad spectrum herbicides for broadleaf weed control such as glyphosate and 2,4-D. In 2023, soybean growers in Eaton County, Michigan, reported repeated failures to control *A. artemisiifolia* populations in non-GM soybean using two PPO-inhibiting herbicides, sulfentrazone and fomesafen (Supplementary Figure 1). In response, seed samples from two suspected resistant populations (Ambel61 and Ambel62) were submitted to the Michigan State University Plant & Pest Diagnostics laboratory to investigate the underlying mechanisms of resistance. Resistance was first characterized through greenhouse dose–response assays, followed by molecular analyses to identify potential genetic alterations associated with reduced herbicide efficacy. In this study, we report for the first time, the natural occurrence of the novel target-site mutation, Arg-98-Gln (R98Q), in the *PPX2* gene of *A. artemisiifolia* populations collected from Michigan soybean fields and characterize this mutation in terms of potential cross-resistance to other PPO-inhibiting herbicides.

## 2. MATERIAL AND METHODS

### 2.1 Plant growth and dose-response experiments

*A. artemisiifolia* seeds from two populations (Ambel61 and Ambel62, hereafter referred to as R1 and R2) were collected in 2023 from soybean fields in Eaton County, Michigan, where resistance to PPO-inhibiting herbicides was suspected. Seeds from a susceptible population (Ambel2021, hereafter referred to as S) were collected in 2021 at the Michigan State University Agronomy Farm and used as a reference control in this study (Supplementary Figure 1).

Seeds were stored in mesh bags and buried outdoors for 12 weeks during winter to break dormancy and synchronize germination. After dormancy breaking and conditioning, seeds were pre-germinated, transplanted into individual pots (4.5 inches) filled with SureMix commercial potting mix (manufactured by Michigan Grower Products Inc., Galesburg, MI, USA), and grown under greenhouse-controlled conditions (27/21 °C, 16/8 h light/dark) until reaching the 4-6 leaf stage, when herbicide applications were performed. Plants were irrigated daily and fertilized every other day with a commercial fertilizer solution (Osmocote®, ICL Specialty Fertilizers, Dublin, OH, USA) at a concentration of 200 ppm.

Herbicides were applied using a research track sprayer (DeVries 174 Manufacturing, INC^TM^, US) equipped with a single flat-fan spray nozzle (8001EVS, TeeJet Technologies, Wheaton, IL, US) delivering 187 L ha^−1^ at 207 kPa in a single pass. Herbicide treatments were applied POST using commercial formulations at labeled field rates, specifically sulfentrazone (Spartan 4F) at 420 g ai ha ¹ and fomesafen (FlexStar) at 350 g ai ha ¹. For sulfentrazone, doses ranged from 1/8× to 16× the labeled field rate, whereas for fomesafen, doses ranged from 1/8× to 4×. To enhance herbicide efficacy, a non-ionic surfactant (NIS) was added at 0.25% (v/v) according to manufacturer recommendations.

### 2.2 Data analysis

Each herbicide rate was applied to a minimum of five plants per population, and each dose– response experiment was independently repeated in time. Fourteen days after herbicide application, plant survival was visually assessed using a binary scale, where 0 indicated complete plant death and 1 indicated plant survival or regrowth (Supplementary Figure 2). LD_50_ values (herbicide dose required to kill 50% of the plants) and ED_50_ values (herbicide dose required to reduce plant biomass by 50%) were estimated from the dose-response data. Resistance levels (R/S) were estimated as the ratio of the LD_50_ and ED_50_ of the resistant population relative to that of the susceptible population.

Dose-response analyses were performed in R using the *drc* package. For aboveground biomass, a three-parameter log-logistic model (LL.3) was fitted to the data according to the following equation:

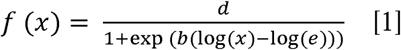

where *d* is the upper asymptote representing the response at zero dose, *b* is the slope at the inflection point, and *e* is the ED_50_. The lower limit was fixed at 0. A joint dose-response model was used including all populations of *A. artemisiifolia* (S, R1 and R2) to allow comparison of response curves among populations. Model fit was assessed using the lack-of-fit test implemented in the modelFit function (*drc* package). The ED_50_ values were estimated for each population from the fitted models. Differences in ED_50_ values among populations were assessed using the EDcomp function in the *drc* package, which performs statistical comparisons of effective doses between curves.

Plant survival data were analyzed using a binomial dose-response model with a two-parameter log-logistic function (LL.2, binomial family):

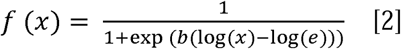

where *b* is the slope at the inflection point and *e* is the LD_50_. The upper and lower limits are fixed at 1 and 0, respectively. As for biomass, a combined model including all populations was used to test for differences among populations. Model fit was assessed using the lack-of-fit test implemented in the modelFit function (*drc* package), and the LD_50_ values were estimated and compared among populations using the EDcomp function.

### 2.3 PPO2 amplification and sequencing

#### 2.3.1 DNA extraction

Leaf tissue from 30 *A. artemisiifolia* plants, including susceptible and survivors from both resistant populations selected across the dose-response range, was collected for genomic DNA extraction. Samples were first flash-frozen in liquid nitrogen and ground using a TissueLyser II system (Qiagen, Hilden Germany). DNA was extracted following the modified cetyltrimethylammonium bromide (CTAB) protocol described by Aboul-Maaty and Oraby, (2019). Briefly, 1000 μl of 1% hot CTAB extraction buffer (65 °C) supplemented with 2% β-mercaptoethanol were added to ground plant tissue and incubated for 1 h at 65 °C with periodic vortexing. Following incubation, samples were centrifuged for 2 min at maximum speed to pellet debris, and 1 μl of RNase (10 mg ml^-1^) was added, mixed gently, and incubated at 37 °C for 30 min. An equal volume (600 μl) of chloroform:isoamyl alcohol (24:1) was added, samples were vortexed until homogenized and then centrifuged at maximum speed for 3 min. The upper aqueous phase was carefully transferred to a new tube, and 400 μl of chloroform:isoamyl alcohol (24:1) were added, followed by vortexing and centrifugation at maximum speed for 3 min. The top layer (400 μl) was removed to a new tube, and DNA precipitation was initiated by adding 1/10 volume (40 μl) of 3 M sodium acetate and 2.0 volumes (800 μl) of cold 100% ethanol. Samples were mixed and centrifuged for 2 min to pellet the DNA. The pellet was washed with 400 μl of cold 70% ethanol, air-dried, and resuspended in 30 μl of DNase/RNase-free water.

DNA concentration and purity were assessed using a NanoDrop spectrophotometer (Thermo Fisher Scientific, Waltham, MA, USA).

#### 2.3.2 Primer design and PCR conditions

To facilitate amplification and sequencing, and due to its length and sequence complexity, the *PPX2* gene was divided into three fragments, each designed to include a single nucleotide polymorphism (SNP) associated with resistance to PPO-inhibiting herbicides (see Table 1 for primer sequences, fragment sizes, and PCR conditions). Polymerase chain reactions (PCR) were conducted in a final volume of 25 µl, containing 5 µl of 5X Q5 Reaction Buffer, 0.5 µl of 10 mM dNTPs, 0.25 µl of Q5 High-Fidelity DNA Polymerase (New England Biolabs, Ipswich, MA, USA), 5 µl of 5X Q5 High GC Enhancer, 1.25 µl each of forward and reverse primers (final concentration of 0.5 µM), 1 µl of DNA template, and 10.75 µl of nuclease-free water. The PCR cycling conditions consisted of an initial denaturation at 98 °C for 2 min, followed by 40 cycles of 98 °C for 30 seconds, annealing at the specific temperature to each primer pair (Table 1) for 30 s, and extension at 72 °C for the corresponding duration (Table 1). A final extension step was performed at 72 °C for 4 min. PCR products were visualized on a 1% agarose gel stained with GelRed nucleic acid stain (Sigma-Aldrich, St. Louis, MO, USA) and purified using the QIAquick PCR Purification Kit (Qiagen, Hilden, Germany) prior to Nanopore sequencing.

**Table 1.**
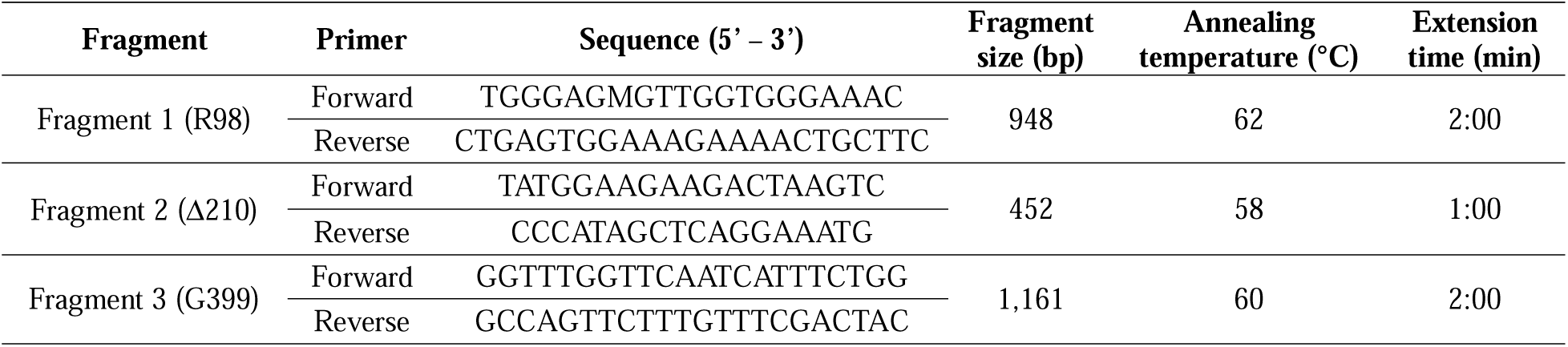
Primer sequences and PCR conditions used for amplification of *PPX2* gene fragments in *A. artemisiifolia*. Fragment 1 includes the R98 single nucleotide polimorfism (SNP), fragment 2 includes the Δ210 SNP and fragment 3 includes the G399 SNP positions. Each fragment was amplified using species-specific forward and reverse primers. The table includes the expected fragment size (in base pairs), annealing temperature (°C), and extension time (minutes) used in PCR amplification for each target region.

### 2.4 Computational modelling

Molecular docking simulations were performed to evaluate the impact of resistance-associated mutations, particularly the Arg-98-Leu (R98L) and the Arg-98-Gln (R98Q) substitutions, on herbicide binding to the PPO2 enzyme. Three-dimensional (3D) structures of the PPO2 protein were predicted using the AI-based tool AlphaFold3, based on the amino acid sequence obtained from the genome annotation provided by the International Weed Genomics Consortium (https://www.weedgenomics.org/)^14^. Both the R98L and R98Q mutations identified in resistant populations were introduced into the protein sequences, and both wild type and mutant variants were independently modeled. The resulting protein structures were subsequently processed in UCSF Chimera v1.19 for structural optimization and energy minimization, including the addition of hydrogen atoms, removal of water molecules, and correction of steric clashes prior to docking. The chemical structures of sulfentrazone, fomesafen, lactofen, saflufenacil, and flumioxazin were retrieved from PubChem in SDF format. Ligands were converted from SDF to PDB format and 3D conformations were generated using Open Babel v3.1.0. Both ligand and protein structures were then converted to PDBQT format using Open Babel v3.1.0 to prepare them for docking. Docking simulations were carried out using AutoDock Vina v1.1.2 ^15^. The docking grid box was centered at x = −2.81, y = 0.32, and z = 4.46, with dimensions of 20 × 20 × 20 Å, covering the predicted PPO2 binding pocket. Simulations were conducted with an exhaustiveness of 8, generating up to 9 binding modes per ligand within an energy range of 3 kcal mol ¹. All parameters were kept consistent across simulations to ensure comparability between wild type and mutant PPO2 structures. Docking poses and protein-ligand interactions were analyzed and visualized using PyMOL v2.5.4 (Schrödinger, LLC).

### 2.5 *In vitro* inhibition studies of PPO2

Complete description of the expression and purification of PPO variant proteins and the enzymatic assay to determine protein activity and median inhibitory concentration (IC_50_) is provided in Rangani et al. (2019).

## 3. RESULTS AND DISCUSSION

### 3.1 Herbicide dose-response assays to quantify resistance levels in *Ambrosia artemisiifolia*

The susceptible (S) *A. artemisiifolia* population was found to have an LD_50_ value of 0.37× the field rate of sulfentrazone (420 g ai ha□¹), whereas the resistant populations R1 and R2 showed significantly higher LD□□ values of 13.6× and 9.1×, respectively (Figure 1A, E), indicating a substantial resistant phenotype when sulfentrazone is applied POST. Resistance levels (R/S), calculated as the ratio of the LD□□ of the resistant populations to that of the susceptible population, were 36-fold and 24-fold for R1 and R2, respectively. Similarly, biomass reduction analysis showed ED_50_ values of 0.06× for the S population and 0.85× and 0.51× for R1 and R2, corresponding to R/S ratios of 14-fold and 8.5-fold, respectively (Figure 1C; Table 2).

**Figure 1.**
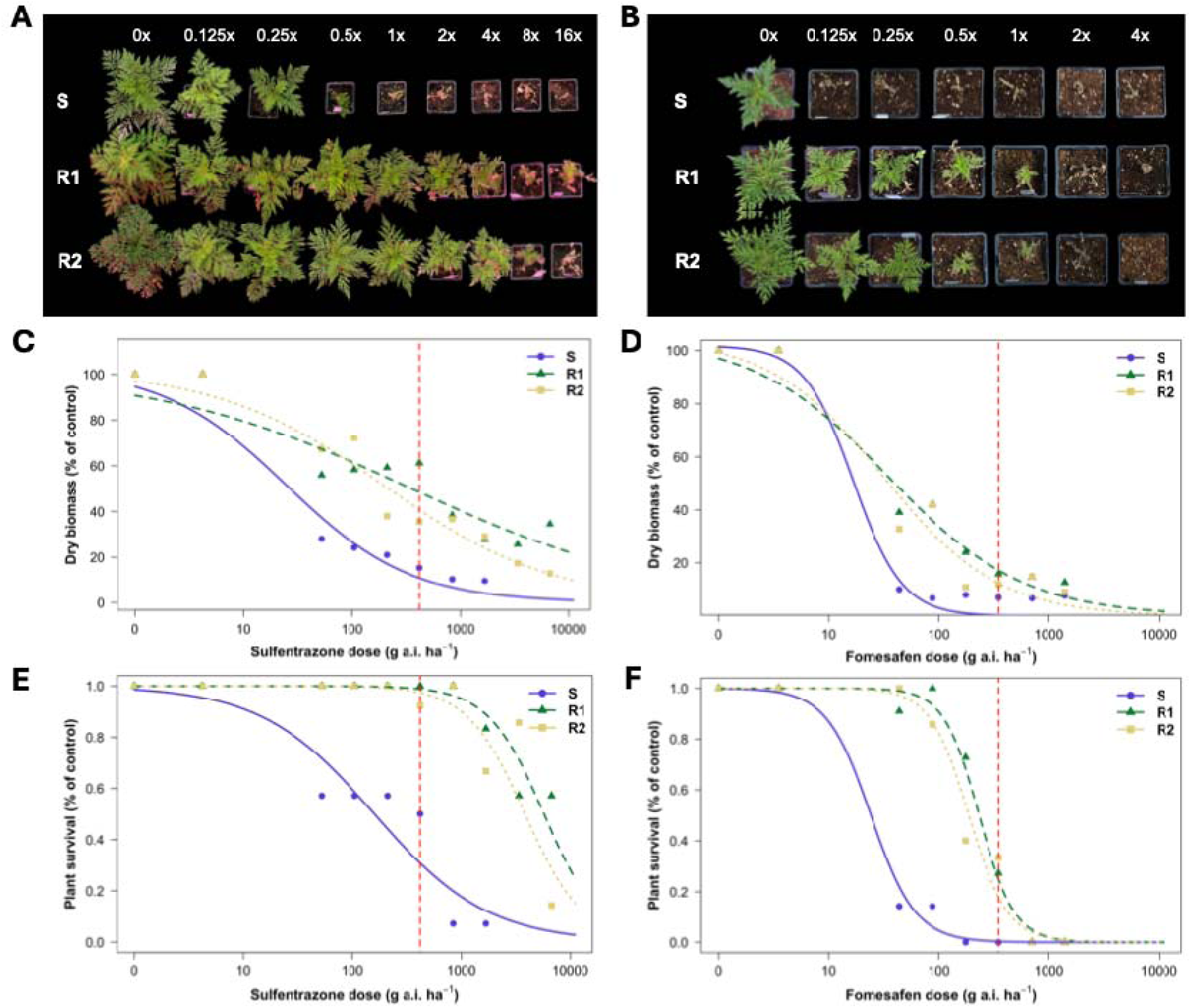
Dose–response curves of *A. artemisiifolia* populations to PPO-inhibiting herbicides applied under post-emergence conditions: (a) sulfentrazone (Spartan 4F) and (b) fomesafen (FlexStar). Herbicide rates on the x-axis are expressed as multiples of the labeled field rate (×), corresponding to 420 g ai ha ¹ for sulfentrazone and 350 g ai ha ¹ for fomesafen. Dry biomass (c, d) and plant survival (e, f) were calculated by using the Equation 1 and Equation 2 displayed in the text, respectively, and are expressed as a percentage of the untreated control. In all plots, the susceptible population (S) is shown in blue, while the resistant populations R1 and R2 are shown in green and yellow, respectively. The red dashed vertical line indicates the labeled field rate for each herbicide.

**Table 2.** Estimated LD and ED_50_ values of sulfentrazone and fomesafen for three *A. artemisiifolia* populations: susceptible (S), and resistant (R1 and R2). Field rate, ED_50_ and LD_50_ values are expressed in grams of active ingredient per hectare (g ai ha ¹). ED_50_ corresponds to the herbicide dose required to reduce biomass by 50% and LD corresponds to the herbicide rate required to kill 50% of the plants. Field rate equivalent is express in × times the labeled field rate. Resistance levels (R/S) were calculated as the ratio of the ED_50_ or LD_50_ of the resistant population relative to the susceptible population.

| PPO-inhibiting herbicide | Population | Field rate | ED<sub>50</sub> | Field rate equivalent (×) | R/S | LD<sub>50</sub> | Field rate equivalent (×) | R/S |
| --- | --- | --- | --- | --- | --- | --- | --- | --- |
| Sulfentrazone | S | 420 | 27<sup>b</sup> | 0.06 | / | 159<sup>b</sup> | 0.37 | / |
|  | R1 |  | 353<sup>a</sup> | 0.85 | 14 | 5729<sup>a</sup> | 13.64 | 36 |
|  | R2 |  | 216<sup>a</sup> | 0.51 | 8.5 | 3840<sup>a</sup> | 9.14 | 24 |
| Fomesafen | S | 350 | 16<sup>b</sup> | 0.04 | / | 24<sup>b</sup> | 0.07 | / |
|  | R1 |  | 34<sup>a</sup> | 0.09 | 2.2 | 232<sup>a</sup> | 0.66 | 9.1 |
|  | R2 |  | 35<sup>a</sup> | 0.10 | 2.5 | 187<sup>a</sup> | 0.53 | 7.6 |

For fomesafen, the LD_50_ value for the S population was 0.07× the labelled field rate (350 g ai ha ¹), while R1 and R2 exhibited significantly higher LD_50_ values of 0.66× and 0.53×, corresponding to R/S ratios of 9.1-fold and 7.6-fold, respectively (Figure 1B, F; Table 2). Similarly, ED_50_ values for biomass reduction were 0.04× for the susceptible population, and 0.09× and 0.10× for R1 and R2, respectively (Figure 1D; Table 2). Although these results indicate a shift in sensitivity relative to the susceptible population, all values remained below the labeled field rate, suggesting reduced sensitivity rather than confirmed resistance to fomesafen.

Rousonelos et al. (2012) also observed a range of resistance levels in *A. artemisiifolia* populations when PPO herbicides were applied POST, ranging from 3- to 80-fold. The differential response between sulfentrazone and fomesafen observed in our populations reflects the complex nature of PPO inhibitor resistance patterns, which can vary among herbicides within this mode of action group, as previously reported in other species such as *A. palmeri* or *E. heterophylla* ^9,16,17^.

### 3.2 Identification of PPO2 target-site mutations in *A. artemisiifolia* populations

Nanopore sequencing was performed on 30 plants (10 survivor plants from R1, 10 from R2, and 10 susceptible plants from S) selected across the dose-response range to screen for single-nucleotide polymorphisms (SNPs) at codons 98, 210, and 399 of the *PPX2* gene, which have been previously associated with target-site resistance to PPO-inhibiting herbicides ^11,18^ (Table 1). Analysis of the three amplified *PPX2* gene fragments revealed substitutions at codon 98 in individuals from the resistant populations R1 and R2. Specifically, we found two substitutions in the first position, where guanine (G) was replaced by thymine (T) resulting in an arginine-to-leucine (R98L), and guanine (G) to adenine (A), resulting in arginine-to-glutamine (R98Q) substitution (Figure 2). Among the 10 individuals sequenced from the R1 population, three plants carried the R98L substitution only, three carried the R98Q substitution only, and three retained the wild-type sequence at codon 98. In addition, one individual was heterozygous and carried both resistance-associated variants at codon 98. In the R2 population, the two substitutions occurred at similar frequencies, with four plants carrying R98L, four carrying R98Q, and two retaining the wild-type sequence. Neither the R98L nor the R98Q substitution was detected in any of the 10 individuals sequenced from the susceptible population. Notably, this R98Q substitution represents the first report of this specific mutation in field collected *A. artemisiifolia* populations. Furthermore, no resistance-associated mutations were detected at codon 210 or codon 399 in any of the individuals screened (Figure 2).

**Figure 2.**
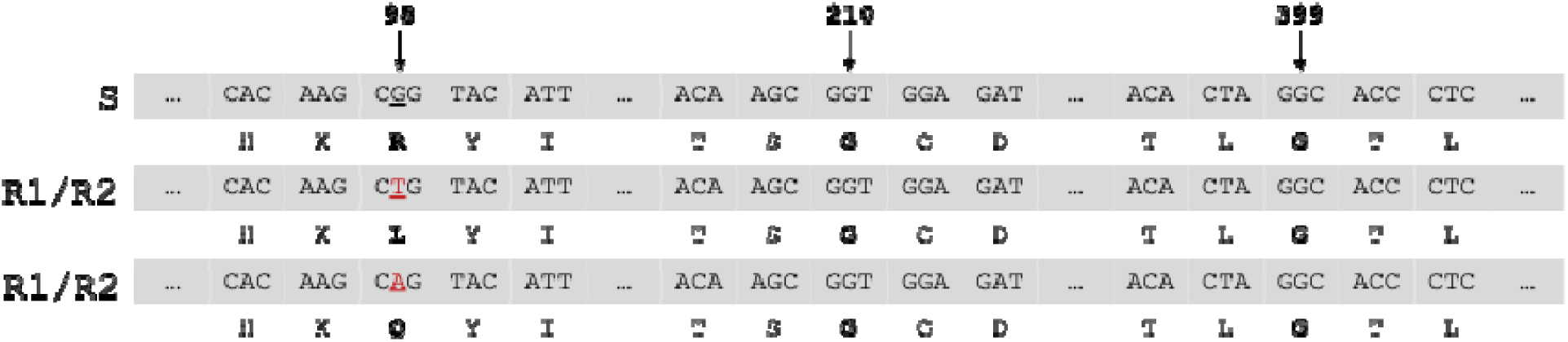
Graphical representation of the single nucleotide polymorphisms (SNPs) identified in the *PPX2* gene of three *A. artemisiifolia* populations collected from Michigan soybean fields (S, R1, and R2). The nucleotide sequence is shown on the top line, with the corresponding amino acid sequence shown directly below. SNP positions (98, 210 and 399) are highlighted in bold, and the nucleotide substitutions are indicated in red and underlined.

Rousonelos et al. (2012) already characterized PPO inhibitor resistance in *A. artemisiifolia* populations from Delaware (USA), identifying the R98L mutation in the *PPX2* gene, which conferred resistance to acifluorfen. This mutation has also been confirmed in giant ragweed (*Ambrosia trifida* L.) populations, establishing the R98 position as a critical site for PPO inhibitor binding across *Ambrosia* species^19^. The absence of mutations at codon 210 or codon 399 in suggests that resistance mechanisms in these *A. artemisiifolia* populations are focused on residue 98, unlike other species such as *A. palmeri* where the ΔG210 deletion and G399A substitutions are among the key resistance mechanisms^7,11,12^. Additionally, the presence of resistant individuals lacking mutations at codon 98 suggests that additional mechanisms beyond PPO2 target-site alterations may contribute to resistance in these populations. Similar combinations of target-site and non-target-site resistance mechanisms have been reported in other species, including *A. palmeri*, highlighting the complex nature of resistance evolution to PPO-inhibiting herbicides^20^.

### 3.3 Effects of the R98Q substitution in PPO2 on herbicide binding

To further explore the potential structural basis of resistance, *in silico* molecular docking simulations were performed between the PPO2 protein and several PPO-inhibiting herbicides from different chemical families, including sulfentrazone, fomesafen, lactofen, saflufenacil, and flumioxazin (Figure 3). Docking simulations were conducted using both the wild type PPO2 protein and the mutant variant carrying both the R98L and R98Q substitutions observed in our field-populations. Binding affinities (kcal mol ¹) corresponding to the best-scoring docking pose predicted by AutoDock Vina for each protein–ligand pair are presented in Table 3, where more negative binding affinity values indicate stronger predicted protein–ligand interaction^15^.

**Figure 3.**
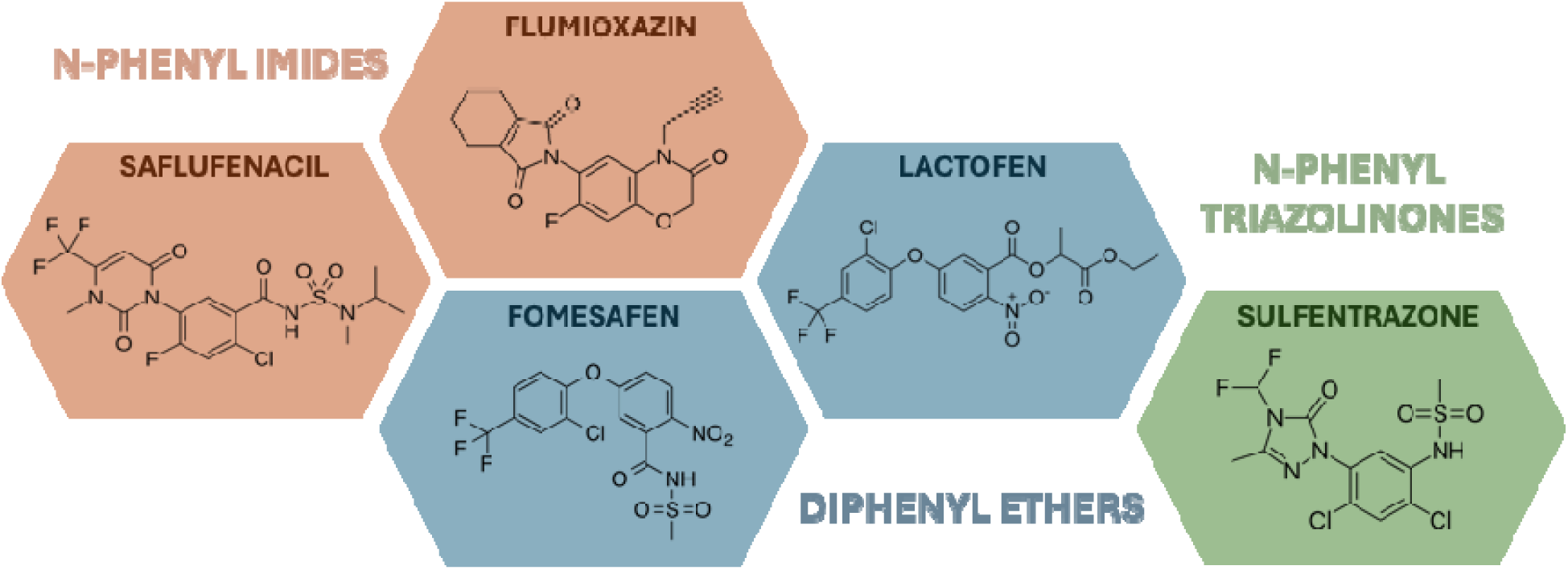
Chemical structures of selected PPO-inhibiting herbicides used in this study and representing major chemical families, including N-phenyl imides in orange (saflufenacil and flumioxazin), diphenyl ethers in blue (fomesafen and lactofen), and N-phenyl triazolinones in green (sulfentrazone).

**Table 3.** Predicted binding affinities (kcal mol ¹) obtained from molecular docking simulations between PPO2 and five PPO-inhibiting herbicides (sulfentrazone, fomesafen, lactofen, saflufenacil, and flumioxazin). Values correspond to the best-scoring docking pose predicted by AutoDock Vina for each protein–ligand pair. Simulations were performed using PPO2 models representing the wild type sequence and both the R98L and R98Q mutant variants of *A. artemisiifolia*. More negative values indicate stronger predicted binding.

| PPO-inhibiting herbicide | Binding affinities (kcal mol<sup>-1</sup>) |  |  |
| --- | --- | --- | --- |
|  | Wild type | R98Q | R98L |
| Sulfentrazone | -7.0 | -5.8 | -5.9 |
| Fomesafen | -8.6 | -6.2 | -7.0 |
| Lactofen | -8.0 | -6.0 | -6.4 |
| Saflufenacil | -7.5 | -6.4 | -6.8 |
| Flumioxazin | -8.1 | -6.3 | -7.0 |

As shown in Table 3, both the R98L and R98Q substitutions resulted in lower predicted binding affinities for all herbicides tested when compared to wild type PPO2, including sulfentrazone (– 5.8 kcal mol□¹ and –5.9 kcal mol□¹ vs –7.0 kcal mol□¹), fomesafen (–6.2 kcal mol□¹ and –7.0 kcal mol□¹ vs –8.6 kcal mol□¹), lactofen (–6.0 kcal mol□¹ and –6.4 kcal mol□¹ vs –8.0 kcal mol□¹), saflufenacil (–6.4 kcal mol□¹ and –6.8 kcal mol□¹ vs –7.5 kcal mol□¹), and flumioxazin (–6.3 kcal mol ¹ and –7.0 kcal mol□¹ vs -8.1 kcal mol□¹), suggesting reduced herbicide binding in the mutated models. These computational predictions are consistent with experimental findings from *A. trifida*, where the R98L substitution similarly diminished docking affinity for lactofen and fomesafen^19^. Notably, the R98Q mutation is predicted to decrease binding to all five PPO herbicides to a greater degree than the previously reported R98L mutation, indicating it may confer stronger resistance.

A visual representation of the molecular docking output between the sulfentrazone and the PPO2 predicted proteins for both wild type and R98Q is shown in Figure 4. The binding pocket, defined as residues within 4 Å of the ligand, comprised 20 amino acids in the wild type PPO2 model, including both hydrophobic (e.g., Val169, Phe344, Leu347) and polar residues (e.g., Asn67, Thr68, Ser174). Notably, Arg98 was located within the pocket, suggesting a potential role in ligand stabilization through electrostatic or hydrogen-bond interactions (Figure 4, left). Upon introduction of the R98Q substitution, the number of residues within 4 Å of the ligand decreased to 10 in the mutant PPO2, indicating a reorganization of the local binding environment. In the R98Q model, the contribution of Arg98 is lost and replaced by a neutral polar residue Gln98, which is not included in the binding pocket under the same distance criterion (Figure 4, right). This structural change was accompanied by a reduction in predicted binding affinity, from –7.0 kcal mol ¹ in the wild type PPO2 to –5.8 kcal mol ¹ in the R98Q variant, consistent with weaker ligand stabilization in the mutated protein. The substitution of a positively charged arginine with a neutral glutamine likely reduces charge-assisted hydrogen bonding and alters the geometry of the binding site, resulting in less favorable interactions and increased solvent exposure within the pocket.

**Figure 4.**
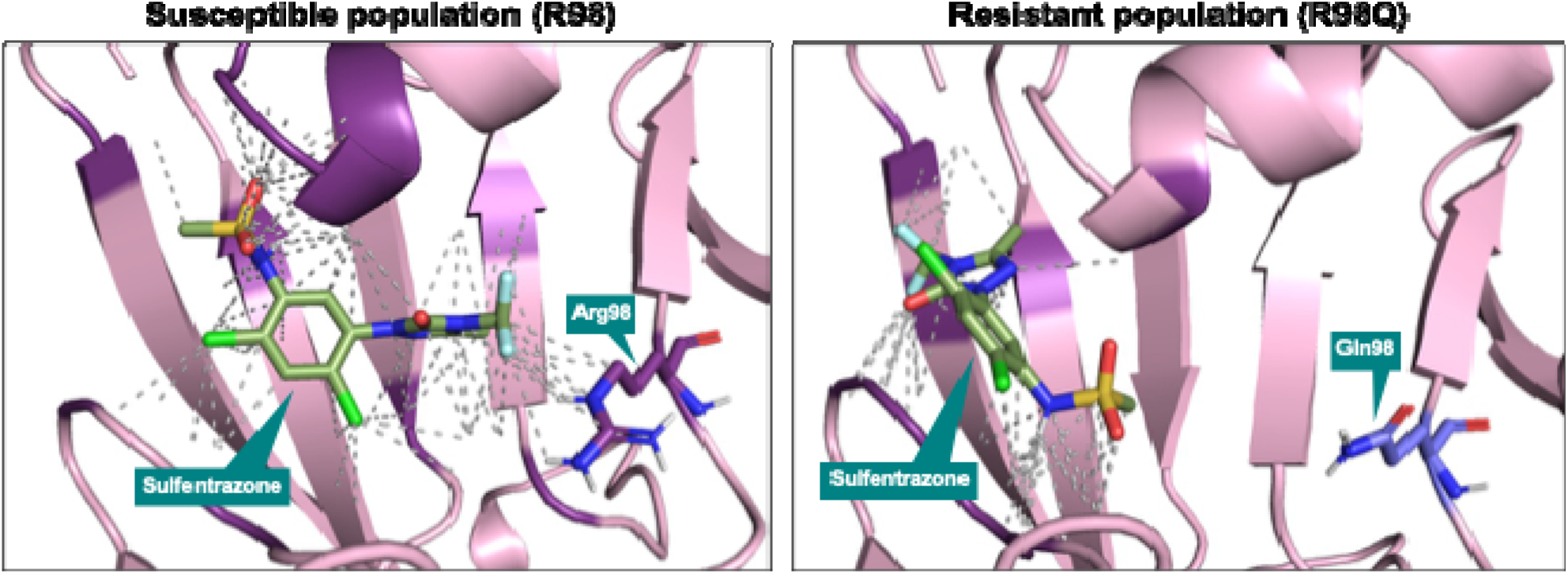
Molecular docking models of wild type and mutant PPO2 proteins in complex with sulfentrazone. The PPO2 protein structure is shown in pink, while the docked sulfentrazone molecule is represented in green. The herbicide-binding pocket is highlighted in violet and defined as residues within 4 Å of the ligand. Interactions between sulfentrazone and the binding-site environment within this distance cutoff are indicated by grey dashed lines. Residue 98 is highlighted in violet for Arg98 (wild type) and purple for Gln98 (mutant), illustrating the position and structural impact of the R98Q substitution. Docking simulations were performed using AutoDock Vina, and protein structures were predicted with AlphaFold3.

This mechanism is consistent with previous experimental evidence showing that substitutions at this position can disrupt salt bridge formation and increase solvent accessibility, thereby reducing herbicide-target affinity^21^. The importance of Arg98 in modulating PPO inhibitor binding has also been highlighted in structural and computational studies of PPO enzymes, which demonstrate that this residue plays a key role in positioning the tetrapyrrole substrate via ionic interactions within the active site^22,23^. Furthermore, computational approaches have been widely used to predict the impact of amino acid substitutions on herbicide binding affinity, including PPO resistance mechanisms in related species^19,24^, supporting the validity of the docking-based interpretation presented here.

### 3.4 Inhibition of R98Q by PPO inhibiting herbicides

The inhibitory activity of multiple PPO-inhibiting herbicides was evaluated against wild type PPO2 and the R98Q PPO2 variant, *in vitro* to validate the binding prediction from predictive molecular docking. Herbicide sensitivity was expressed as IC_50_ values [M], where M represents molarity (mol L^-1^), and resistance was quantified using the resistance index (RI), calculated as the ratio of the IC value for the R98Q variant relative to wild type PPO2 (Table 4). Herbicides with IC_50_ values greater than 1.00 × 10^-5^ M were considered highly resistant under the assay conditions, indicating a high level of enzyme insensitivity.

**Table 4.** Effects of protoporphyrinogen oxidase (PPO) inhibitors on *in vitro* enzyme activity of *A. artemisiifolia* PPO2 wild type enzyme and R98Q resistant enzyme variant. Herbicide sensitivity was expressed as IC_50_ values [M], where M represents molarity (mol L^-1^), and resistance was quantified using the resistance index (RI), calculated as the ratio of the IC value for the R98Q variant relative to wild type PPO2.

| PPO chemical family | Herbicides | Wild type sensitivity (IC<sub>50</sub>) [M] | R98Q sensitivity (IC<sub>50</sub>) [M] | Resistance index |
| --- | --- | --- | --- | --- |
| Diphenyl ether | Fomesafen | 1.12E-06 | 4.09E-07 | 0 |
|  | Acifluorfen | 3.50E-06 | 6.37E-07 | 0 |
|  | Lactofen | 5.23E-09 | 6.18E-09 | 1 |
|  | Nitrofen | 2.50E-07 | 1.35E-07 | 1 |
| N-Phenylimides | Butafenacil | 5.97E-10 | 6.08E-08 | 102 |
|  | Trifludimoxazin | 4.98E-10 | 6.29E-08 | 126 |
|  | Tiafenacil | 4.60E-09 | 1.58E-06 | 343 |
|  | Epyrifenacil | 6.21E-10 | 1.47E-06 | 2367 |
|  | Saflufenacil | 1.84E-07 | >0.00001 | ND |
| N-Phenyltriazolinones | Carfentrazone-ethyl | 6.83E-09 | 5.10E-06 | 747 |
|  | Sulfentrazone | 1.81E-06 | >0.00001 | ND |
| Other | Pyraflufen-ethyl | 7.52E-10 | 5.78E-08 | 77 |
|  | Oxadiazon | 8.58E-09 | 1.95E-06 | 227 |
|  | Pyraclonil | 1.66E-08 | 1.12E-05 | 675 |

Wild type PPO2 was strongly inhibited by all tested herbicides, with IC_50_ values ranging from the micromolar to sub-nanomolar scale, demonstrating broad susceptibility across chemical classes. In contrast, the novel R98Q substitution resulted in marked and compound-specific reductions in inhibition. Diphenyl ether herbicides (fomesafen, lactofen, nitrofen, and acifluorfen) retained comparable activity against wild type and R98Q. IC_50_ values for the R98Q variant remained similar to those obtained for wild type, resulting in RI values of 0 or 1, indicating that the R98Q mutation does not confer resistance to this chemical class *in vitro*. These results highlight the value of combining computational and experimental approaches, as docking simulations may not fully reproduce the dynamic interactions involved in herbicide binding. In contrast, a shift in IC_50_ was observed for several N-phenylimide herbicides. Butafenacil and trifludimoxazin exhibited increases in IC_50_ values for the R98Q variant compared with wild type enzyme, with RI values of 102 and 126, respectively. However, both butafenacil and trifludimoxazin inhibit the R98Q enzyme in the nM range, suggesting that while a shift in sensitivity could be observed, both herbicides still strongly inhibit the target mutant enzyme. A greater shift in IC_50_ was observed for tiafenacil (RI = 343) and epyrifenacil (RI = 2367). For saflufenacil, the IC_50_ toward the R98Q enzyme was >1.00 × 10 M, indicating pronounced enzyme insensitivity and preventing RI determination. Importantly, the reduced sensitivity to fomesafen observed in whole-plant experiments was not fully reflected in the *in vitro* enzyme assays, suggesting that additional mechanisms may contribute to the resistance phenotype for this herbicide.

Among N-phenyltriazolinone herbicides, carfentrazone-ethyl and sulfentrazone showed a strong loss of inhibitory activity against the R98Q variant. Carfentrazone-ethyl exhibited a substantial increase in IC_50_ value, corresponding to an RI of 747, whereas sulfentrazone displayed an IC50 > 1.00 × 10 M, indicating a high level of resistance. These patterns of resistance to triazolinone herbicides are consistent with previous findings demonstrating that mutations at homologous positions in related species confer resistance to this chemical class, with the R128L mutation (equivalent to R98L in *A. artemisiifolia*) causing resistance to triazolinones among other chemical families^12^. The enzyme-level resistance observed for sulfentrazone correlates well with our molecular docking predictions showing reduced binding affinity for this herbicide in the R98Q variant, and with whole-plant dose-response experiments conducted in the greenhouse, where field populations exhibited significant resistance to sulfentrazone. These findings are also consistent with previous functional studies in heterologous expression systems, where the equivalent R128Q substitution (corresponding to R98Q in *A. artemisiifolia*) altered PPO2 sensitivity in engineered *E. coli* systems ^21^. In that study, the R128Q mutation was evaluated in combination with the ΔG210 deletion, rather than as a single substitution, and was associated with different levels of enzyme activity depending on the herbicide tested. Although these results were obtained in a double-mutant background and in a heterologous system, they nevertheless support the functional relevance of residue 98/128 in PPO inhibitor binding and enzyme sensitivity.

The remaining herbicides, classified as ‘others’, also displayed reduced inhibition of the R98Q variant. Pyraflufen showed a large shift in IC_50_ value relative to wild type (RI = 675), while pyraflufen-ethyl exhibited a moderate resistance level (RI = 77). Oxadiazon likewise demonstrated a clear resistance phenotype, with an RI value of 227. Overall, these data demonstrate that the R98Q mutation in PPO2 confers strong and selective resistance to multiple PPO-inhibiting herbicides, particularly in some N-phenylimides and N-phenyltriazolinones, while diphenyl ether herbicides remain effective. In some cases, such as for sulfentrazone, the IC_50_ related to the R98Q variant exceeded the maximum tested concentration, highlighting a high degree of enzyme insensitivity associated with this substitution.

## 4. CONCLUSION

This study represents the first natural occurrence of the arginine-to-glutamine (R98Q) substitution at position 98 of PPO2 in field-collected *A. artemisiifolia* populations from Michigan. This mutation was associated with resistance to sulfentrazone and reduced sensitivity to fomesafen in whole-plant assays, indicating that changes at residue 98 can significantly impact PPO inhibitor efficacy. These results highlight the rapid evolution and diversification of target-site resistance mechanisms in *A. artemisiifolia* and underscore the importance of continued monitoring of PPO-inhibiting herbicide performance. Importantly, first natural occurrence of this novel substitution also highlights the need for the development of rapid diagnostic tools to enable early detection and tracking of emerging resistance mutations in weed populations. Further research is needed to determine whether additional mechanisms beyond PPO2 target-site substitutions contribute to the observed variability in herbicide response in these populations.

## ACKNOWLEDGEMENTS

The authors would like to acknowledge the undergraduate students at Michigan State University for their assistance with greenhouse experiments and sample processing. Funding for this project was provided by both the Michigan Soybean Committee and the Herbicide Resistance Monitoring Network [HERMON] project funded by the United Soybean Board (Project #26- 206-S-C-2-G). Additional Support was provided by Project GREEEN (Project #GR24-077) for protein folding and docking.

## CONFLICTS OF INTEREST

AP and JL are employees of BASF Company (Germany). The remaining authors declare that the research was carried out without any commercial or financial relationships that could be considered a conflict of interest.

