## Supplemental Material for "Characterization of a novel R98Q mutation that confers resistance to sulfentrazone in common ragweed (*Ambrosia artemisiifolia*) populations from Michigan"

**Supplementary Material**

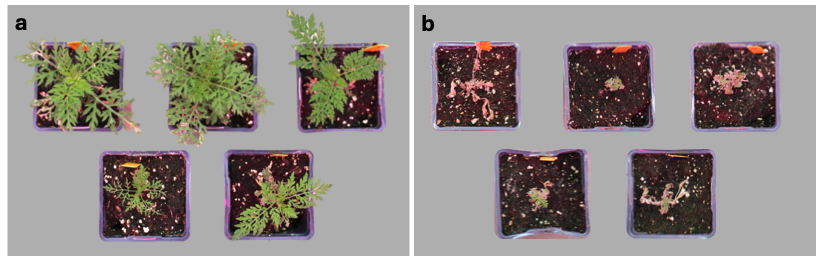

Supplementary Figure S1. Visual criteria used to assess plant survival following herbicide application in greenhouse dose–response and frequency assays. Plants were evaluated 14 days after treatment and classified as alive (1) or dead (0) based on visual assessment. (a) Representative examples of plants scored as alive (1), showing survival and/or significant regrowth. (b) Representative examples of plants scored as dead (0), showing complete or almost complete plant necrosis.
